# Site-Specific Investigation of DNA Holliday Junction Dynamics and Structure with 6-Methylisoxanthopterin, a Fluorescent Guanine Analog

**DOI:** 10.1101/2024.04.19.590264

**Authors:** Zane Lombardo, Ishita Mukerji

**Affiliations:** Department of Molecular Biology and Biochemistry, Molecular Biophysics Program, Wesleyan University, 52 Lawn Ave, Middletown, Connecticut 06459, United States

**Keywords:** 6-MI, 6-Methylisoxanthopterin, Holliday junction, fluorescence, fluorescent guanine analog, DNA dynamics

## Abstract

DNA Holliday Junction (HJ) formation and resolution is requisite for maintaining genomic stability in processes such as replication fork reversal and double-strand break repair. If HJs are not resolved, chromosome disjunction and aneuploidy result, hallmarks of tumor cells. To understand the structural features that lead to processing of these four-stranded joint molecule structures, we seek to identify structural and dynamic features unique to the central junction core. We incorporate the fluorescent guanine analog 6-methylisoxanthopterin (6-MI) at ten different locations throughout a model HJ structure to obtain site-specific information regarding the structure and dynamics of bases relative to those in a comparable sequence context in duplex DNA. These comparisons were accomplished through measuring fluorescence lifetime, relative brightness, fluorescence anisotropy, and thermodynamic stability, along with fluorescence quenching assays. These time-resolved and steady-state fluorescence measurements demonstrate that the structural distortions imposed by strand crossing result in increased solvent exposure, less stacking of bases and greater extrahelical nature of bases within the junction core. The 6-MI base analogs in the junction reflect these structural changes through an increase in intensity relative to those in the duplex. Molecular dynamics simulations performed using a model HJ indicate the primary sources of deformation are in the shift and twist parameters of the bases at the central junction step. These results suggest that junction-binding proteins may use the unique structure and dynamics of the bases at the core for recognition.

## Introduction

DNA Holliday junctions (HJs) are branched molecules consisting of four double stranded arms and are essential intermediates for processes like replication fork reversal, and double-strand break repair. Proper DNA Holliday junction formation and resolution must occur to maintain genomic stability during these important processes. Improper resolution of HJs can result in chromosome disjunction and aneuploidy leading to inherited genetic disorders and cancer. Since 1964 when Robin Holliday initially suggested the existence of this intermediate central to homologous recombination [1] there have been many studies investigating the overall structure and dynamics of the branched DNA molecule [2, 3] and how different proteins interact to resolve them [4, 5]. Single-molecule fluorescence and other studies have shown HJs primarily exist in two stacked anti-parallel conformational isomers, iso-I or iso-II, or an unstacked open-X conformation, with the stacked X isomers being favored in the presence of mono- and polyvalent cations [6-8]. The well-characterized J3 junction used in this study adopts the iso-II conformation 80% of the time where the strands designated B and R are the exchanging strands and the H and X strands are the continuous strands [9-13] (Fig 1A). J3 is an HJ that is base pair matched with no homology rendering it immobile or incapable of branch migration.

**Figure 1.**
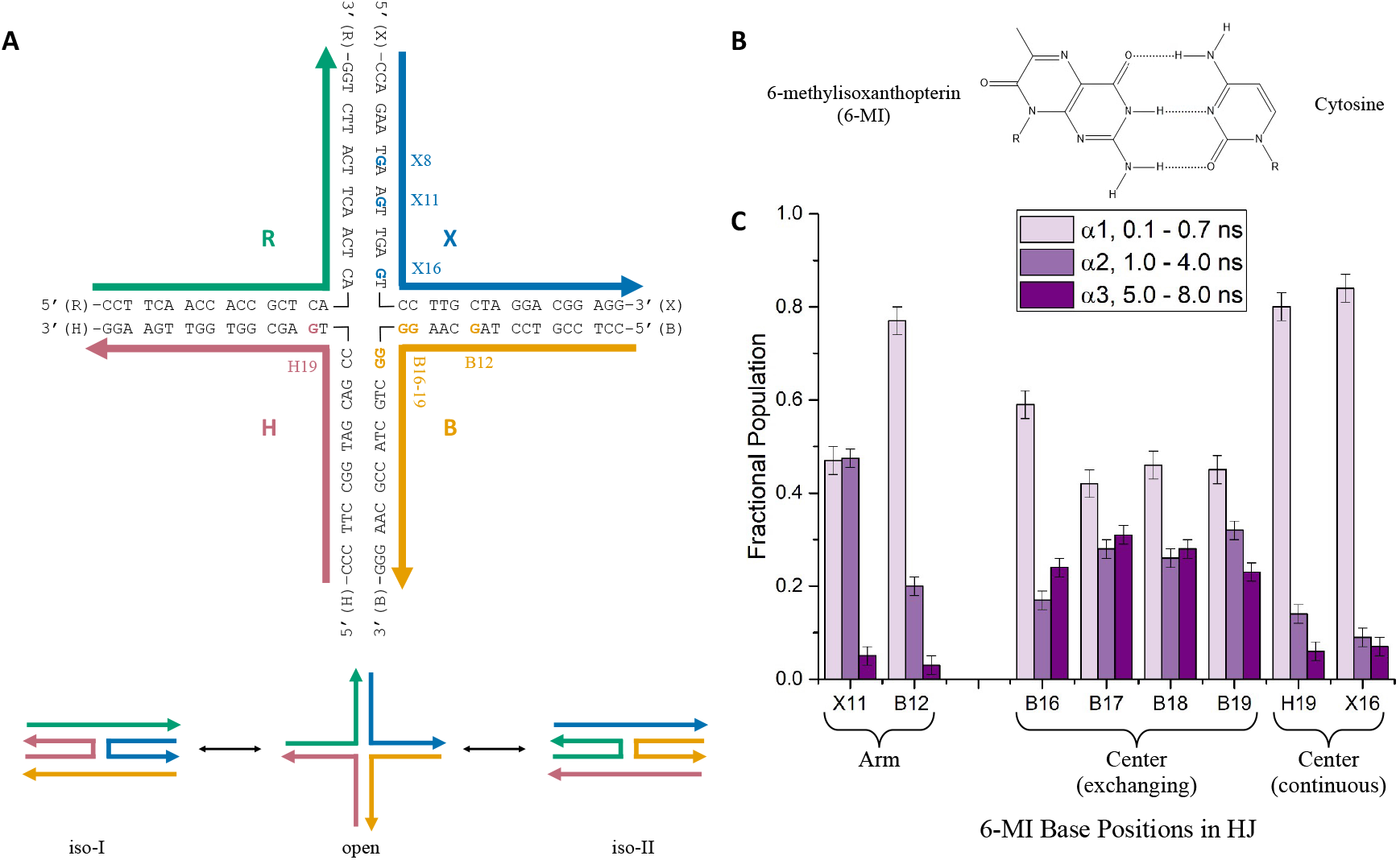
6-MI containing J3 Holliday junctions. (A) A schematic representation showing the sequence of the J3 Holliday junction and probe locations (bold and shown in strand color). In our studies, each junction only has one probe. J3 is made up of four strands: B (yellow), H (red), R (green), and X (blue). The J3 Holliday junction is detected in three possible conformations: iso-I, open, and iso-II. Iso-II is the preferred state (80%) under the conditions used in this study. (B) The chemical structure of the fluorescent guanine analog 6-MI, which forms Watson-Crick hydrogen bonds with cytosine. (C) Fractional populations of the lifetime components are obtained from fitting of fluorescence lifetime decays to a sum of exponentials as described in the text. Fractional populations of mid- and long-range components increase in the junction center (B16-19) compared to the arms (X11, B12). Samples contained 200 nM DNA in a 10 mM Tris, pH 7.5, 100 mM NaCl and 5 mM MgCl_2_ buffer. Fit parameters are given in Supporting Information: Table S2.

Amongst the abundance of studies that have examined the overall structure and conformer distribution of HJs, there has been little investigation of the local structure and dynamics of DNA bases at the core of the HJ. Prior NMR studies have shown that bases at the junction center maintain Watson-Crick type base pairing but deviate from a regular B-DNA conformation [14-16]. Importantly, the identity of bases at the central core are critical for determining HJ conformer distribution and dynamics [17] and they are the site for many protein-HJ interactions [18-22]. Thus, understanding the structure and dynamics of the central bases is important for establishing junction behavior and how these repair and recombination intermediates interact with proteins.

To further determine the local structure and dynamics of DNA bases in Holliday junctions, we employed a fluorescent DNA base analog, 6-methylisoxanthopterin (6-MI), to report on the local environment and motions at the level of a single DNA base. This fluorescent guanine analog forms Watson-Crick H-bonds with cytosine in double-stranded DNA while minimally perturbing normal structure (Fig. 1B) [23-25]. Previous studies have demonstrated that 6-MI fluorescence is sensitive to local environment within DNA, is a useful tool for measuring the local motions of DNA bases in different sequence or structural contexts and is a sensitive reporter of protein binding [23, 26-28]. Previously, 6-MI revealed significant differences in the local motion of DNA base mismatches, which could be related to the ability of the mismatch repair protein Msh2-Msh6 to recognize specific mismatch types [29]. The probe has also been successfully used as a molecular sensor for G-quadruplex formation where an increase in fluorescence intensity, resulting from a reduction in base stacking interactions, signifies G-quadruplex formation [30, 31]. In this work, we use 6-MI to understand how the structure and dynamics of bases in different locations within the junction, particularly the junction center, leads to specific recognition by junction-binding proteins. We further examine if the base local environment within the junction differs from duplex DNA in the same sequence context.

To accomplish this, we have incorporated 6-MI at ten different locations throughout our HJ model system to gather site specific information. Fluorescence lifetime measurements, the relative brightness and fluorescence anisotropy at each location, along with fluorescence quenching assays demonstrate that structural distortions imposed by strand exchange results in increased solvent exposure, reduced stacking interactions between bases, and greater extrahelicity at the junction core. Molecular dynamics simulations using a model HJ further suggest that the source of deformation is primarily in the shift and twist parameters of the bases at the junction center and that bases at the HJ center in the exchanging strands have increased motion. These results which give both base-specific and site-specific information demonstrate the utility of 6-MI for probing nucleic acid structure and dynamics. They further suggest that protein recognition of junctions may be driven by the non-canonical, dynamic structure of the exchanging strand bases at the junction center.

## Materials and Methods

### Sample Preparation

6-MI containing oligonucleotides were purchased from Fidelity Oligos LLC (Gaithersburg, MD) in PAGE-purified form. HPLC purified complementary strands were purchased from Integrated DNA Technologies (Coralville, IA). All other reagents were purchased from Sigma-Aldrich (St. Louis, MO) unless otherwise indicated. Single stranded DNA was stored at -20 °C prior to use. Duplex and HJ DNA were prepared by preparing equal molar amounts of the 6-MI-containing and complementary strands in a 10 mM Tris, pH 8.0 buffer containing 300 mM NaCl and 1 mM EDTA, heating at 75 ºC for 6 hrs followed by slow cooling to room temperature in a water bath.

### Steady State Fluorescence Measurements

The fluorescence intensities of 6-MI containing J3 samples relative to duplex DNA were measured on a Horiba Fluoromax-4 spectrofluorometer (Edison, NJ) at 10 ºC at 430 nm. Intensities were determined from fluorescence emission spectra using an excitation wavelength of 340 nm. The fluorescence emission was scanned from 400 to 500 nm at a resolution of 1 nm/pt with an integration of 0.5s/pt with an excitation and emission bandpass of 2 nm. Ratios of peak intensity values were determined from the spectrum of each J3 substrate divided by the peak intensity values obtained from the spectrum of the equivalent duplex. Fluorescence anisotropy measurements were made with excitation and emission wavelengths of 340 nm and 430 nm, respectively. The excitation and emission slits had a bandpass of 5 nm. Fluorescence anisotropy ratios were determined in the same manner as intensity ratios. All values were determined from at least three different experiments for each J3 or duplex substrate and errors are reported as the standard deviation. Each DNA substrate was prepared in Buffer A (10 mM Tris, pH 7.5, 100 mM NaCl, and 5 mM MgCl_2_) to a final concentration of 200 nM.

### Fluorescence Quenching Assays

KI was titrated from 0 to 160 mM into a 400 nM solution of 6-MI-containing duplex or HJ DNA. The ionic strength of the solution was held constant at 200 mM K^+^ by balancing the KI and the KCl concentrations in a 10 mM Tris, pH 7.5 buffer. Samples were excited at 340 nm and background corrected emission was collected at 430 nm with a 5 second integration and excitation and emission slits had a 3 nm bandpass. Samples were maintained at constant temperature of 10 °C.

The quencher accessible fraction was determined from non-linear curve-fitting of the data using a modified Stern-Volmer equation [32]:

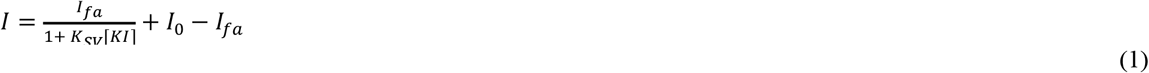

Where *I* represents the intensity, *I*_0_ represents the initial intensity, *I*_*f*a_ is the intensity of the quencher accessible fraction, and *K*_*SV*_ is the Stern-Volmer constant. The quencher accessible fraction (*f*_*a*_) was calculated from the average of three separate quenching experiments and the error is reported as the standard deviation.

### Time-Resolved Fluorescence Lifetime Measurements

Time-correlated single-photon counting (TCSPC) was performed using a Photon Technology International TimeMaster instrument. Samples containing 100-300 nM DNA were prepared in Buffer A and excited with a 375 nm pulsed picosecond diode laser with a repetition rate of 1 MHz (Becker & Hickl). Fluorescence emission was detected at 460 nm with emission slits set at a 20 nm bandpass using a 450 nm cutoff filter and an emission polarizer set to 54.7°. All samples were stirred and maintained at a constant temperature of 10 °C over the course of the measurement. Integrity of the samples was verified post irradiation through native gel electrophoresis [29]. Decays were collected over a 55 ns time range with 4096 channels until a maximum of 20000 counts were obtained in the peak channel. The instrument response function was collected by measuring scattered light from a colloidal suspension of Ludox (Sigma Aldrich). The lifetime decays were fit to a sum of exponentials using the following equation:

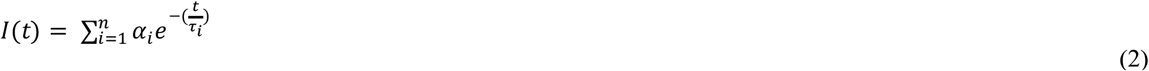

where *I(t)* is the intensity at time *t*, and *α* is the amplitude of the ith component and represents the fractional population with a lifetime of τ_*i*_. Fitting and analysis were performed using either Globals Unlimited [33] or FelixGX (Photon Technologies International) software. Goodness of fit was determined through an examination of residuals and reduction of chi-squared values to 0.9-1.3. DEF substrates were fit to a sum of two exponentials and all other substrates were fit to a sum of three exponentials. The intensity-weighted mean fluorescence lifetimes (τ_*f*_) and relative amplitudes *α*_*i*_ were calculated from the fits and are averaged from three independent experiments using the following expressions:

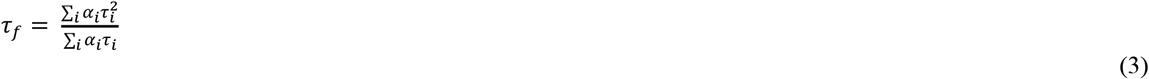

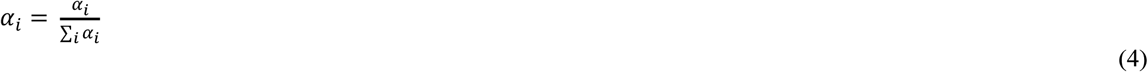

### Molecular Dynamics Setup and Analysis

The starting Holliday junction structure was obtained from Prof. Wilma Olson (Rutgers University) [34]. Molecular dynamics simulations for the J3 Holliday junction were performed using the GPU PMEMD version of AMBER 16.0 and AMBER 18.0 suite of programs with the parmbsc1 forcefield for the DNA [35-37]. The system was solvated using a TIP3P [38] water model in a 12.0 Å octahedral box with Na^+^ or K^+^ ions to a final concentration of 150 mM NaCl or KCl. The particle mesh Ewald algorithm [39, 40] and a 10 Å Lennard-Jones cutoff were used to treat long-range electrostatic interactions. Initial calculations were energy minimized with four separate minimizations of 1000 steepest descent (SD) and 500 conjugate gradient (CG) steps with harmonic constraints of 100, 100, 10, 0 kcal/mol on solute and 20, 0, 0, 0 kcal/mol on the ions, respectively. This was followed by slow heating to 300 K, and four separate equilibrations of 10 ps steps with harmonic constraints of 25, 25, 15, 5 kcal/mol on solute and 20, 20, 10, 0 kcal/mol on the ions respectively. The production steps were performed in conjunction with NPT ensemble conditions and Berendsen algorithm [41]. The SHAKE [42] constraints were applied to all the bonds including hydrogen bonds with an integration time of 2 fs and the trajectory snapshots were saved every 2 ps until the final MD simulation time was reached.

Stability and convergence of the simulations were assessed with root mean-square deviations (RMSD) with respect to the average structure. The 3DNA webserver was used to calculate the helical parameters for the average structure obtained from the MD simulations [43]. Root mean square fluctuations (RMSF) by residue were obtained using the CPPTRAJ RMSF command with time frames of 200 ps using the average structure as reference [44]. The RMSF was calculated using a sliding window of three bases as a means for estimating the local motion of the middle base with respect to its adjacent neighbors. The center of mass for each nucleotide base was determined using PyMol [45]. Distances between the center of mass for adjacent bases were measured using the average structure. The B-DNA reference was generated by analyzing a homoduplex DNA crystal structure (PDB: 1BNA) in the same manner [46].

## Results

To understand the structural features that lead to protein recognition and processing of the four-stranded branched DNA molecules known as four-way or Holliday junctions (HJ), we compare the environment of DNA bases within the arms and center of HJs relative to duplex DNA. To ensure that we maintained the same sequence context, 34 bp 6-MI-containing strands were annealed with corresponding complementary strands to form either duplex or HJ DNA (Table S1). We used ten different 6-MI probe positions to observe single base dynamics at distinct locations in the well-characterized J3 Holliday junction. The J3 junction exists primarily in the iso-II conformation (80%) and we use this conformation for designating the continuous and exchanging strands.

Nevertheless, we note that 20% of the population is in the iso-I conformation, implying that any effects observed are likely stronger than detected [7-13]. Herein, we identify specific positions by indicating the junction strand followed by the position from the 5’ end of that strand. For example, X8 indicates the 6-MI probe is in the X strand and at base position 8 from the 5’ end of that strand. Residue positions, 8, 11, and 12 are located in a junction arm at positions where we expect the DNA structure to be most like canonical B-DNA (Fig. 1A) [18, 47]. Positions 16 through 19 in the B strand of J3 are all guanines and can be readily substituted with 6-MI giving us access to bases at the junction center (positions 17 and 18) and to bases located one position from the center (positions 16 and 19) on an exchanging strand without changing the DNA sequence. In addition, position 19 on the H strand and position 16 on the X strand are guanines and are used to probe positions one base from the center on continuous strands (Fig. 1A). By using these positions, we were able to incorporate 6-MI at various locations within the junction without altering the sequence.

### How does the environment of the HJ center compare with the HJ arms?

To probe the environment of the different locations within the HJs, we employed time-resolved fluorescence spectroscopy and measured the lifetimes of 6-MI probes located in the arms and at the center of HJs. Fluorescence lifetime decays were well-described by a sum of exponentials in which three lifetime components, short (0.1-0.7 ns), medium (1-4 ns), and long (5-8 ns), were needed to fit the decays as observed previously (Supporting Information: Fig. S1). The short lifetime component arises from 6-MI stacking with adjacent bases, the long lifetime component is attributed to an extrahelical conformation of 6-MI, while the medium lifetime component is assigned to an intermediate conformation [23, 24, 48, 49]. (Fig.1 C). The amplitude of each decay component corresponds to the fractional population of that component. We use these fractional populations to compare the differences in probe environment in the different junction locations. When the 6-MI probe is incorporated into the junction arms, the fractional population of the long component is greatly reduced, and the majority of the decay is described by the short and mid-range lifetime components. This is consistent with a probe environment where the fluorescence is mainly quenched through stacking and collisional interactions with adjacent bases. In contrast, when 6-MI is incorporated at the HJ center, the fractional populations are more evenly distributed between the three lifetime components. This equality in distribution is mainly caused by an increase in population of the long lifetime component and a decrease in population of the short component (Fig. 1C). This re-distribution of populations is primarily observed for the probes in the exchanging strands (B16-B19). In the case of probes on the continuous strand (H19 and X16), this effect is less pronounced, and the fractional populations of the long-lived components only increase slightly. As the junction exhibits an 80:20 population distribution for the iso-II and iso-I conformations [7-13] the H19 and X16 probes will be in an exchanging strand for a fraction of the time, possibly leading to the longer lifetimes observed. The increase in population of the long lifetime component probably arises from a conformation in which 6-MI is experiencing less quenching from collisional and stacking interactions with neighboring bases by visiting an extrahelical, solvent exposed state more frequently. At the HJ center, bases on the exchanging strands experience more torsional strain when in the stacked-X conformation and are likely to adopt an extrahelical conformation to relieve the strain as discussed below.

We employed the previously characterized duplex-enhanced fluorescence (DEF) sequence ATFAA [23, 24] (F = 6-MI) to observe this effect with greater sensitivity. We incorporated the DEF sequence in either the arms (X8) or center (X17) of the HJ (Fig 2). The X8 position serves as a control since it is in the same sequence context that has been extensively characterized in duplex DNA [24]. As the 6-MI is in the arm of the junction where the helical parameters are expected to resemble duplex DNA, we anticipated that probe dynamics would be similar to those previously determined. In our earlier study, we found that the ATFAA sequence stabilized the 6-MI probe and reduced collisional quenching, resulting in an increase in fluorescence intensity upon duplex formation [24]. In contrast, when 6-MI is placed into the X8 arm position, we see an approximately 20% decrease in fluorescence intensity (Fig 2B) relative to the duplex control. We attribute this intensity decrease to a disruption of the DNA structure that rigidly holds the probe in place and keeps it from interacting with neighboring bases. When 6-MI is placed in the HJ center within the ATFAA sequence we detect an approximately 35% decrease in fluorescence intensity compared to homoduplex DNA (Fig 2B). This larger decrease relative to that observed in the arms is suggestive of a greater loss of rigidity in the B-DNA structure, consistent with more collisional quenching and increased motion of the probe. While 6-MI in the DEF sequence experiences some loss of structural stability in the HJ arm, the even larger decrease in fluorescence intensity observed for X17 at the center is suggestive of greater flexibility in the structure of bases at the HJ center relative to the arm.

**Figure 2.**
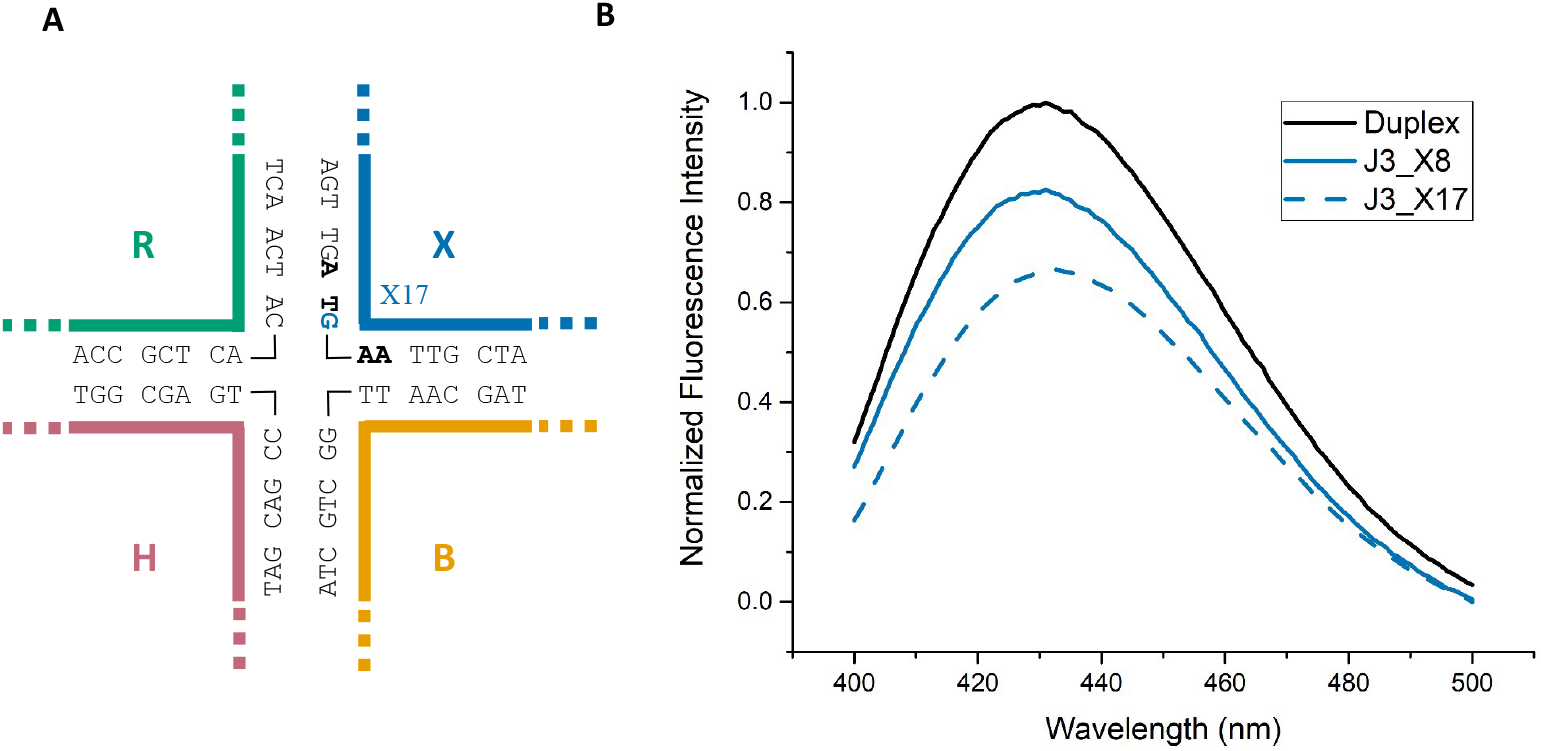
Duplex-enhanced fluorescence sequences in duplex or HJ DNA. (A) A schematic representation of the J3 junction sequence modified to incorporate the ATFAA sequence at the HJ core (Blue G is replaced with 6-MI). (B) 6-MI in the ATFAA sequence is brightest in homoduplex DNA (HD, black). Incorporation of the ATFAA sequence into the arm of J3 (X8, blue) results in a roughly 20% decrease in fluorescence intensity. 6-MI is further quenched when the ATFAA sequence is placed at the center of J3 (X17, blue and dashed). All samples were 200 nM DNA in a 10 mM Tris, pH 7.5, 100 mM NaCl and 5 mM MgCl_2_ buffer.

### Molecular dynamics simulations reveal structural perturbations and increased dynamics at HJ center

We developed and performed Molecular Dynamics simulation on a J3 junction model to gain detailed structural information at an atomic level to better interpret our fluorescence results. To analyze the average structure resulting from the MD trajectories, we used the DNA structural analysis tool, 3DNA [43, 50]. Through this analysis, we compared the helical parameters of the junction bases with canonical B-form DNA. Twist and shift base pair step parameters calculated from 3DNA are shown for a 1μs simulation performed in NaCl (Fig. 3A) and a 100 ns simulation in KCl (Supporting Information: Fig. S2). In both the NaCl and KCl simulations, the center junction bases exhibit substantial deviations from the average simulated structure. The core base step bridging positions 17 and 18 on either side of the junction center exhibit sharp changes in structure in which the twist changes by as much as 20 degrees and the shift by ± 1 Å relative to canonical values. As the bases get farther away from the center, the base pair steps more closely resemble B-DNA in the twist and shift helical parameters. We found that analysis of both simulations yielded similar trends suggesting that the identity of the ion did not significantly impact the results.

**Figure 3.**
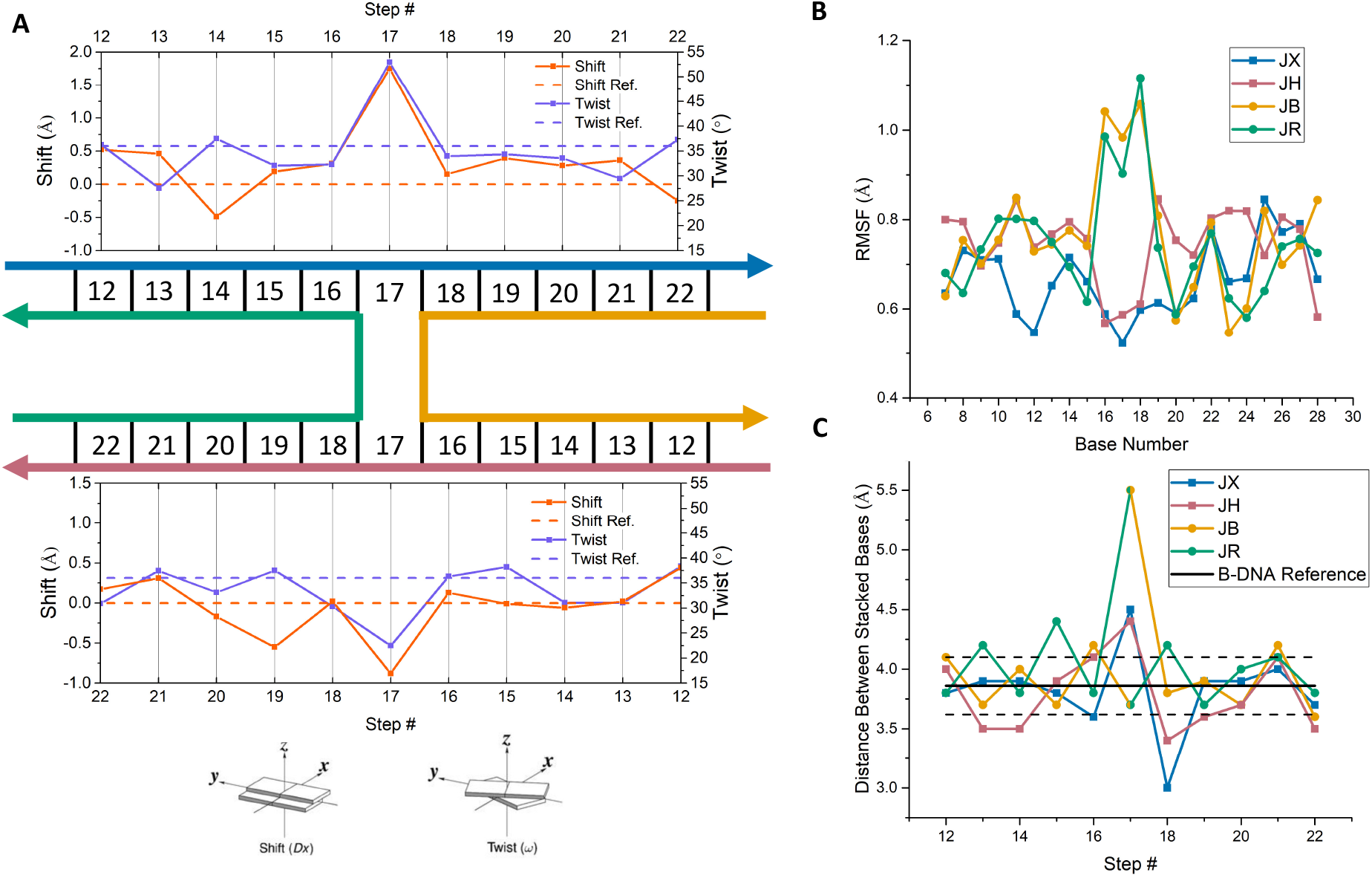
Molecular dynamics simulations of the J3 HJ reveal structural perturbations and increased dynamics at HJ centers. (A) Analyses of a 1 μs simulation were performed using the 3DNA webserver [43] and revealed deviations in the twist (purple) and shift (orange) base pair step parameters of J3 from canonical B-form DNA. Canonical B-DNA values are shown in a dashed line and J3 parameters are depicted with a solid line. Step 17 at the HJ center exhibits the largest deviations. (B) Root-mean-square fluctuations (RMSF) throughout the course of the simulation are shown for bases in the JX (blue), JH (red), JB (yellow) and JR (green) strands. The RMSF values were calculated for individual bases using a sliding window of three bases to estimate the local motion of the middle base of the three bases with respect to its nearest neighbors. The greatest fluctuations are detected for bases at the center of the HJ in the exchanging strands (JB and JR, yellow and green, respectively). (C) Distances between the center of mass for adjacent bases at each base step in the average MD structure. The distance between bases at the HJ center (steps 17, 18) are greater than the average value determined for B-DNA. The standard deviation for the B-DNA reference is shown by the black dashed lines.

Root-mean-square fluctuations (RMSF) were calculated for bases throughout the HJ to determine if base dynamics at the junction center are increased relative to other positions. The RMSF was calculated using AMBER CPPTRAJ [44] and a sliding window of three bases to reduce the contribution of larger global motions of the HJ and focus on local motions of the middle base of the three bases within the context of the nearest neighbors. The terminal bases of the DNA strands were not included because of the common effect of end fraying [51]. Interestingly, the RMSF analysis shows that there is increased motion for positions 16, 17, and 18, but only for the exchanging strands B and R (Fig 3B). In fact, center bases 16, 17, and 18 on the continuous H and X strands seem to be less dynamic on average than bases in the arms of HJs. The increased motion of bases in the exchanging strands observed in this analysis is consistent with the increased population of the extrahelical state observed in our fluorescence lifetime measurements. We estimated stacking interactions by examining the distances between the center of mass of the DNA bases and compared the values with similar measurements performed on standard B-form DNA. These analyses are consistent with those of the shift, twist and RMSF, where bases at the center deviate significantly from standard values (Fig. 3C). Cumulatively, these MD simulation results, which indicate bases at the center are significantly distorted from B-form DNA, are consistent with our spectroscopic results and all together suggest that the DNA bases at the center of a HJ deviate substantially from canonical B-form DNA in structure.

Previous molecular dynamics simulations of HJs consisting of different DNA sequences also observed distortions in the twist and shift parameters for bases at the HJ center and reported that bases in the junction arms have very similar helical parameters to B-form DNA [47, 52, 53]. In addition, coarse-grained simulated melting of J3 shows the center takes the shortest time to melt [54], where the thermostability is inferred from the faster melting of the center relative to the arms and is suggestive of decreased stability in the center. We attribute the lower stability in part to weaker stacking interactions at the HJ center which is correlated with increased lability of those bases.

### Comparison of junction base structure and dynamics with duplex DNA

To examine the environment and dynamics of individual bases in Holliday junctions, we compared the properties of 6-MI probes in HJs and in homoduplex DNA in the same sequence context. Steady state anisotropy, and fluorescence emission spectra were collected for each DNA substrate. This comparison of steady-state anisotropies and fluorescence intensities between junctions and duplexes further indicates increased dynamics of bases at HJ centers when compared to B-form duplex DNA. The ratios of the junction fluorescence intensity to the duplex with the same sequence are shown in Fig. 4A. These intensity ratios clearly show that the 6-MI probe positions at the HJ center (positions 16-19) are more fluorescent than their duplex counterparts. As the probe is moved out of the center and into the arms of the HJ there is less of a difference in intensity which can be seen by looking at the B12, X11, and X8 positions. Comparing the changes in intensity between probes located at the B16-B19 (exchanging) positions to H19 and X16 (continuous) positions also reveals that this effect is greater for bases in the exchanging strand versus the continuous strand (Fig. 4). We interpret our results based on the dominant conformation in solution; however, the conformation distribution is 80:20 and the minor conformation will influence the results and possibly leads to the small changes detected for the H19 and X16 probes. We further observe that 6-MI located in the junction center exhibited the same spectral characteristics as when it is located in a loop or adjacent to a mismatch site [24, 29], consistent with a loss of duplex structure and more frequent excursions to extrahelical or single-stranded states. In general, these results indicate probes at the HJ center are less stacked and experience less collisional quenching from neighboring bases than they would in a duplex DNA.

**Figure 4.**
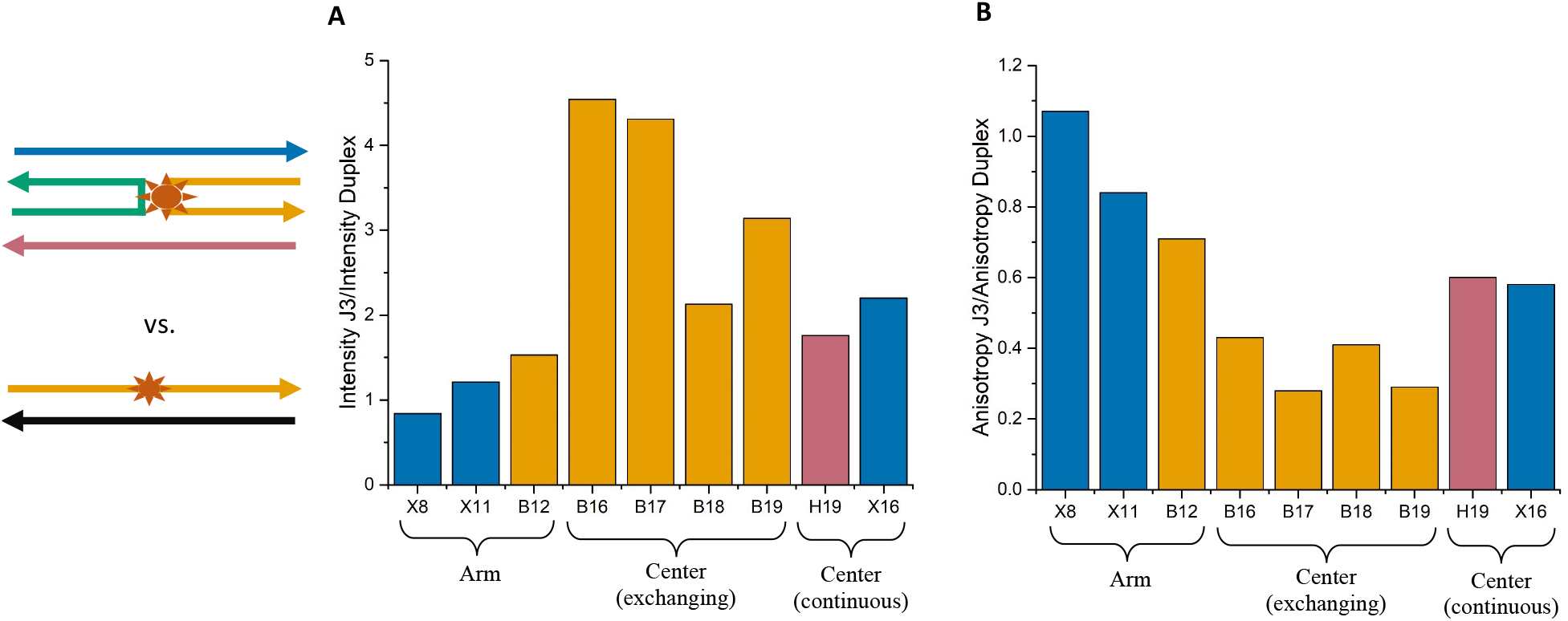
Ratios of fluorescence intensity (A) and anisotropy measurements (B) for 6-MI containing HJs and duplexes in identical sequence contexts. (A) Probes in the HJs exhibit increased fluorescence intensity compared to their duplex counterparts, with the greatest differences observed for locations B16-B19. (B) Fluorescence anisotropy values are lower for 6-MI probes in HJs compared to duplex DNA. This effect is most pronounced for bases at the HJ center in the exchanging strands (B16-B19). Ratios are determined from at least three measurements. Samples were 200 nM DNA in a 10 mM Tris, pH 7.5, 100 mM NaCl and 5 mM MgCl_2_ buffer.

Steady state anisotropy experiments reported on the local and global motions of the 6-MI probe when placed in different locations throughout the molecule and we used this data to explore the differences in the junction itself and compared them to duplex DNA. Our steady state anisotropy measurements (Fig. 4B) indicated the anisotropy of 6-MI in a duplex is higher than 6-MI in the HJ within the same sequence context. It is surprising that the anisotropy of specific locations in the junction were lower than those in the duplex, as the junction is larger and should have a longer global rotation time. As the steady state measurements give a weighted average of local and global motions [32], this finding suggests the local motions in the junction outweigh the global motions of the HJ. As shown in Fig. 4A, 6-MI anisotropy measurements also revealed that this effect is more pronounced for bases located at the center rather than the arms of HJs. We infer from these results that bases contained in the HJ experience greater local dynamics and less quenching from neighboring bases relative to bases in duplex DNA.

Fluorescence lifetime measurements of 6-MI probes in HJ or duplex DNA also point to an increase in local dynamics for bases located in the HJ. We observed an increase in the intensity-weighted fluorescence lifetimes (τ_*f*_) in the HJ compared to duplex DNA (Fig. 5A). This effect is much more pronounced for the B16 probe compared to the B12 probe, consistent with our other measurements that suggested bases at the center are more dynamic and solvent-exposed relative to bases in the arms. The increase in lifetime for probes located in the junction arms further indicated that the helical structure in the arms is less constrained than in the corresponding duplex, suggesting that the torsional stress induced by the center exchanging strands propagates throughout the junction as suggested by our measurements with the duplex-enhanced fluorescence substrates (Fig. 2 and Fig. S3).

**Figure 5.**
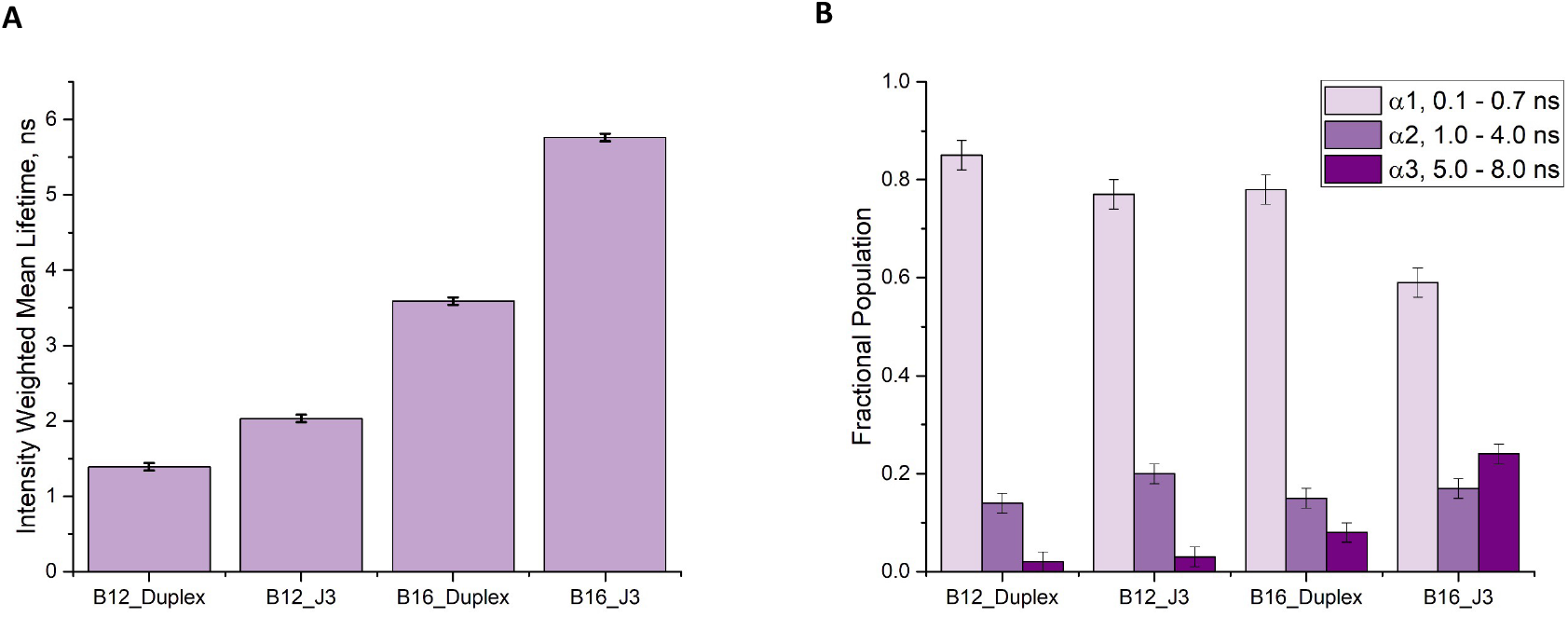
Fluorescence lifetime measurements comparing HJ and duplex DNA for different HJ probe positions. (A) Intensity-weighted fluorescence lifetimes of 6-MI probes in HJs and duplex DNA within the same sequence context were calculated as described in the text and are significantly longer for the B16 probe in the junction. (B) Fractional populations of the lifetime components obtained from analyzing the fluorescence lifetime decays with a sum of exponentials as described in the text. The increase in lifetime at the B16 junction position arises from a shift in fractional population from the shortest lifetime component to the longest. Error bars represent the standard deviation from at least three experiments. Samples were 200 nM DNA in a 10 mM Tris pH 7.5, 100 mM NaCl and 5 mM MgCl_2_ buffer. Fit parameters are given in Supporting Information: Table S3.

We also note that the distribution of fractional populations of the 6-MI fluorescence lifetime components differs between duplex and junction, where an increase in fractional population of the longer-lived components is observed for junction decays (Fig, 5B). The longest lifetime component of the 6-MI probe (5-8 ns) likely arises from a conformation that is more extrahelical in nature and more comparable to that of the monomer dye, while the short component results from collisional quenching interactions with neighboring bases. Analysis of the B12 probe decay demonstrates that the shift in fractional populations mainly occurred between the short (0.1-0.7 ns) and mid-range (1-4 ns) lifetime components. In the case of the center B16 position, the increase in fractional population of the long-lived component was more pronounced and was three times that of the long-lived component measured in the same sequence context in duplex DNA (Fig. 5B), further supporting our finding that bases in the junction center are less stacked and adopt an extrahelical conformation more frequently.

To verify the extrahelical nature of the center bases, we measured the relative solvent exposure of the 6-MI probe in different locations throughout the junction (Fig 6). Quenching of 6-MI fluorescence induced with KI addition provides an estimate of the quencher accessible and inaccessible fraction of the fluorophore. Comparison of data from the X11 and B16 duplexes demonstrates that the quencher accessible fraction depends on sequence context. As shown, in duplex DNA, the B16 position leads to greater quencher accessibility of the 6-MI (73%) relative to the X11 sequence (35%) due to the purine nature of the adjacent bases (Table 1) [26, 55]. Nevertheless, a comparison with the same sequence context in the HJ shows that for the B16 position, quencher accessibility increased by 13% to 86 ± 4% while the probe in the X11 position only experienced half the increase in accessibility, approximately 6% to 41 ± 9% (Table 1) (Fig 6). We note that the difference in quencher accessibility between the duplex and junction arm is within our range of error, suggesting that the environments are comparable. Collectively, all our fluorescence and simulation results point to an environment in the junction center that is more solvent exposed and exhibits greater dynamics than either the junction arms or the corresponding duplexes.

**Table 1.** Quencher accessibility of 6-MI in duplex and HJ substrates determined with KI.

| DNA Substrate<sup>1</sup> | Quencher Accessible Fraction<sup>2</sup> |
| --- | --- |
| B16_Duplex | $0.73 \pm 0.05$ |
| B16_HJ | $0.86 \pm 0.04$ |
| X11_Duplex | $0.35 \pm 0.06$ |
| X11_HJ | $0.41 \pm 0.09$ |

<sup>1</sup>Letter and number indicate location of probe in either duplex or junction. <sup>2</sup>The quencher accessible fraction was determined from non-linear curve-fitting of the data using a modified Stern-Volmer equation (Eq. 3) [32] as described in the text. The quencher accessible fraction ( $f_a$ ) was calculated from the average of three separate quenching experiments and the error is reported as the standard deviation.

**Figure 6.**
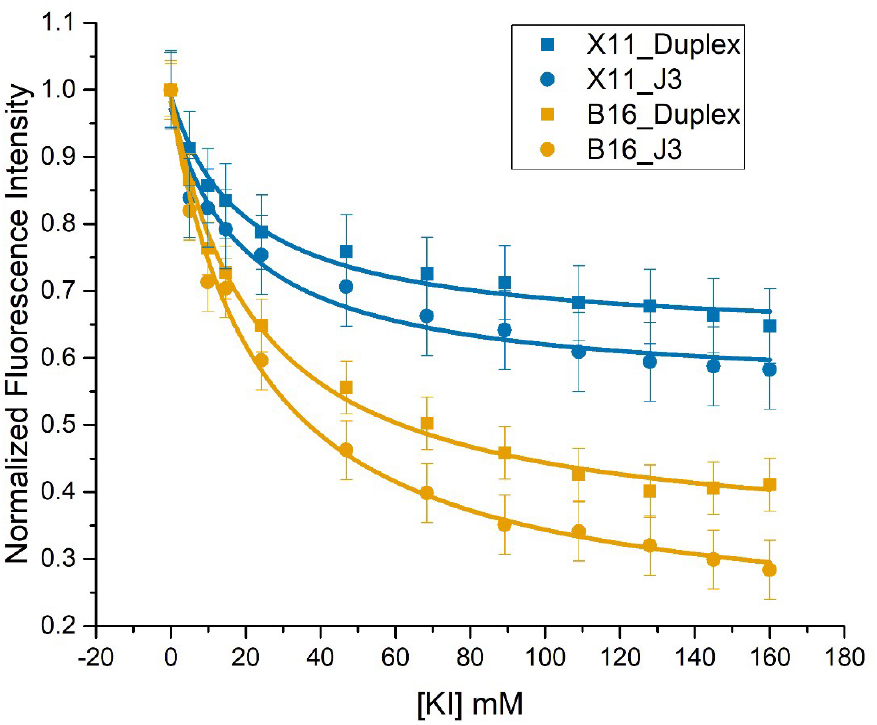
Quencher accessibility of 6-MI in the arms or center of HJs compared to duplex DNA within the same sequence context. The difference in the quencher accessible fraction between HJ and duplex DNA is greater for the position at the center of the HJ (B16) than the arm of the HJ (X11), consistent with the increase in the extrahelical state of 6-MI at the HJ center. The quencher accessible fraction was determined using a modified Stern-Volmer expression as described in the text.

## Discussion

### A Dynamic and Flexible Junction Center

Our time-resolved and steady state fluorescence emission measurements performed at ten unique sites coupled with MD simulations have yielded new information on the local structure and mobility of a 34 bp DNA Holliday junction. This information points to a distorted central junction region in which individual bases adopt extrahelical conformations more often and consequently, experience reduced stacking interactions. Furthermore, our findings show that bases in the junction exchanging strands experience greater perturbations in local structure relative to those in the continuous strands. These unique structural properties of bases at the junction center revealed by 6-MI fluorescence and MD simulation, provide a potential avenue through which proteins both recognize HJs and distinguish the exchanging from the continuous strands.

The current findings are well supported by those obtained with other methods. Through the broadening of proton resonances, NMR studies have also detected greater conformational flexibility in exchanging strands relative to continuous strands [14-16, 56]. A prior study used circular dichroism and differential scanning calorimetry to infer a reduction in base stacking at the junction center [57]. Significantly, our results both confirm and extend these observations as the 6-MI probe is a more sensitive reporter of local structure and, detected reduced base stacking in the arms within a helical turn of the center. Overall, these findings suggest that the junction structure at the central core and extending to proximal bases is more dynamic and less constrained than duplex DNA and possibly presents a likely target for protein recognition and binding, as discussed below.

### Implications for Protein Recognition of Junction Structures

The average structure of our simulated J3 Holliday junction used in this study is depicted in Fig. 7 in comparison with four protein-bound junction structures. Notably, in the protein-junction complexes, the center regions are quite distorted and relatively open with many deviations from canonical B-DNA structure. As the center is a primary point of contact with the proteins, these distortions are likely caused by the protein-junction interaction. Our simulated J3 structure maintains the stacked-X iso-II conformation, and the increased opening at the center is not observed (Fig. 7A). A recent report demonstrated that variations of the standard AMBER forcefields better simulate spontaneous transitions in HJ conformations [58]. Thus, in our case, the AMBER bsc1 forcefield could be over stabilizing the central region through increased stacking interactions; however, we note that we still detect extensive distortions in structure and heightened dynamics (Fig. 3; Fig. S2). Undoubtedly, these distortions would be even more pronounced with the use of the specialized forcefields.

**Figure 7.**
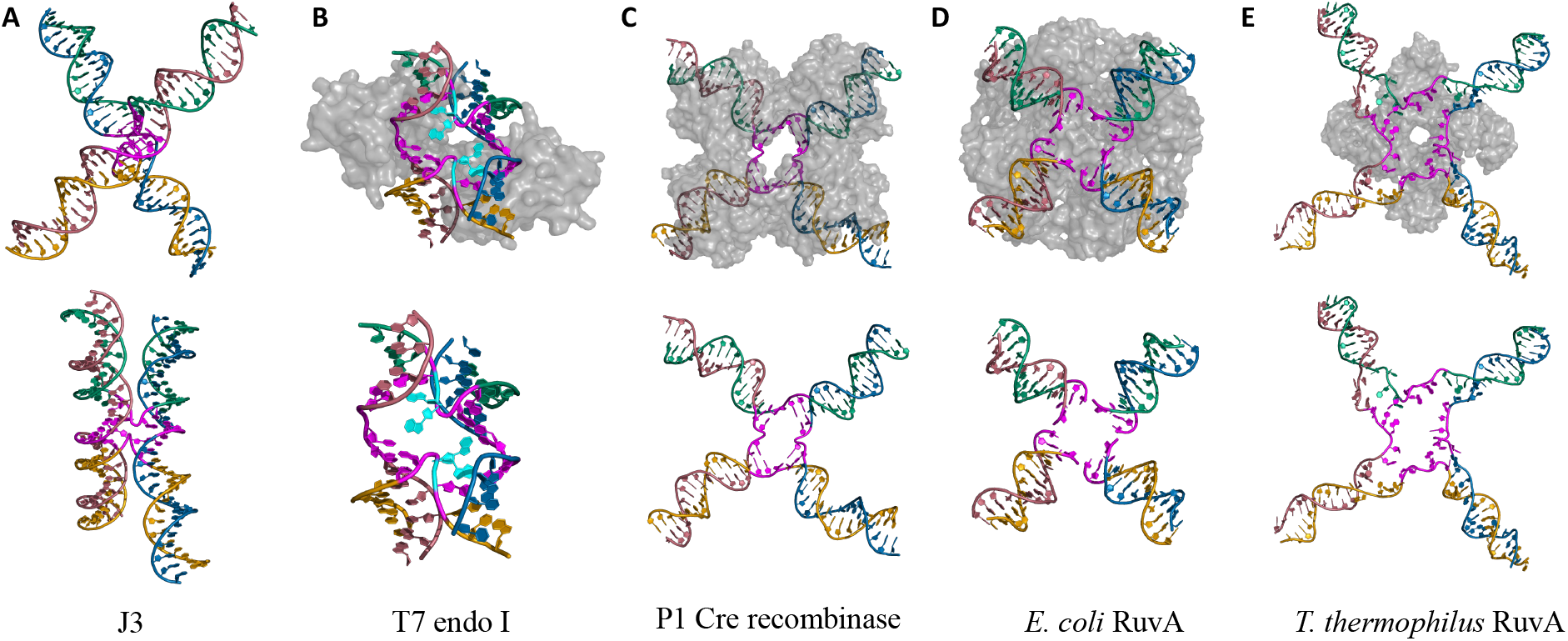
Structural comparison of apo and protein-bound Holliday junctions. Strands in each junction are color coded to match the equivalent arms in the J3 junction used in this study. The four central bases of each strand are highlighted in magenta. (A) The average structure of the apo J3 HJ from 1μs of MD simulation performed in 100 mM NaCl. Conditions for the simulation are given in the text. (B) A T7 endonuclease I (endo I) bound HJ (PDB: 2PFJ) shows bases at the HJ core resolved in two different orientations (highlighted in cyan). (C) P1 Cre recombinase bound HJ (PDB: 2QNC), (D) E. coli RuvA bound HJ (PDB: 1C7Y), and (E) T. thermophilus RuvA bound HJ (PDB: 8GH8) all show significant opening and distortion of the junction center with protein bound.

The loss of base stacking and in some cases base-pairing in the protein-bound structures (Fig. 7B-E) suggests that structural flexibility at the core is an important recognition mechanism for junction-binding proteins [18, 21, 59]. For example, in the T7 endonuclease I bound-HJ structure (Fig. 7B), two bases at the HJ center crystallized in two different conformations (highlighted in cyan). Importantly, this base is located in an exchanging strand on the 5’ side of the core step, making it equivalent to base 17 (B17 or X17) in our study. The presence of two different conformations when bound to protein is consistent with the inherent distortion and flexibility of the B17 and X17 bases we measured in our solution assays. Similarly, the open conformation at the center of the P1 Cre recombinase-HJ complex and larger than average thermal parameters in the crossover region of this structure also point to the increased mobility of this region [59]. The *E. coli* RuvA crystal structure [18] and *T. thermophilus* RuvA EM structure (unpublished, PDB:8GH8) also exhibit opening at the HJ center, which is accompanied by the unstacking and unpairing of bases. Thus, these protein-bound junction structures support and confirm our findings that the junction core represents a labile and dynamic region, in which excursions to extrahelical states potentially facilitate recognition and binding.

In previous work, 6-MI has been shown to be a sensitive reporter of DNA structure either alone or in complex with proteins [23, 24, 26, 27, 29, 60, 61]. Our study further supports and extends these findings as subtle changes in junction structure, undetected by other methods, were reflected through 6-MI fluorescence properties. This sensitivity to junction structure, makes it a useful probe for studying protein-junction interactions.

## Conclusion

In this work, we have used 6-MI to investigate local structure and dynamics of bases in DNA Holliday junctions. Through time-resolved and steady-state fluorescent measurements and molecular dynamics simulations we have shown that the structural distortions imposed by strand crossing result in increased solvent exposure, less stacking of bases and greater extrahelical nature of bases within the junction core. Furthermore, these studies show that bases in the exchanging strands are more labile than those in the continuous strands. These deviations from standard B-form DNA suggest a mechanism through which junction-binding proteins may recognize Holliday junctions and discriminate between strands. The current study further confirms the usefulness of the 6-MI probe for investigating DNA structure and dynamics. As Holliday Junctions are attractive drug targets for cancer therapies [2], we suggest that the sensitivity of 6-MI to junction structure, as demonstrated in this report, makes 6-MI an excellent candidate for developing an effective sensor to screen ligand binding to the junction center.

## Abbreviations

(6-MI): 6-Methylisoxanthopterin
(HJ): Holliday Junction
(TCSPC): Time-correlated single-photon counting
(RMSD): Root mean square deviations
(RMSF): Root mean square fluctuations
(MD): Molecular Dynamics
(NMR): Nuclear Magnetic Resonance
(DEF): duplex-enhanced fluorescence
(AMBER): Assisted Model Building with Energy Refinement

## Acknowledgements

We thank Prof. Wilma Olson (Rutgers University) for the structural coordinates of the model junctions. This work was supported by National Institutes of Health grant R15GM135904 (awarded to IM).

