## Supporting Information for "Site-Specific Investigation of DNA Holliday Junction Dynamics and Structure with 6-Methylisoxanthopterin, a Fluorescent Guanine Analog"

Running title: Site-Specific Investigation of DNA HJ Dynamics Using 6-MI

### Supporting Information (Tables S1-S3; Figures S1-S3)

**Supporting Information: Tables**

Table S1. Sequences of DNA oligomers used in this study. F = 6-MI. The letter denotes the strand of the Holliday junction, while the number represents the position of 6-MI from the 5' end. All strands are 34 bp long

| Name | Sequence |
|---|---|
| JX | 5′ – CCA GAC TGC AGT TGA GTC CTT GCT AGG ACG GAG G – 3′ |
| JX8 | 5′ – CCA GAA T**F**A AGT TGA GTC CTT GCT AGG ACG GAG G – 3′ |
| JX11 | 5′ – CCA GAC TGC A**F**T TGA GTC CTT GCT AGG ACG GAG G – 3′ |
| JX16 | 5′ – CCA GAC TGC AGT TGA **F**TC CTT GCT AGG ACG GAG G – 3′ |
| JX17 | 5′ – CCA GAC TGC AGT TGA T**F**A ATT GCT AGG ACG GAG G – 3′ |
| JB | 5′ – CCT CCG TCC TAG CAA GGG GCT GCT ACC GGA AGG G – 3′ |
| JB12 | 5′ – CCT CCG TCC TA**F** CAA GGG GCT GCT ACC GGA AGG G – 3′ |
| JB16 | 5′ – CCT CCG TCC TAG CAA **F**GG GCT GCT ACC GGA AGG G – 3′ |
| JB17 | 5′ – CCT CCG TCC TAG CAA G**F**G GCT GCT ACC GGA AGG G – 3′ |
| JB18 | 5′ – CCT CCG TCC TAG CAA GG**F** GCT GCT ACC GGA AGG G – 3′ |
| JB19 | 5′ – CCT CCG TCC TAG CAA GGG **F**CT GCT ACC GGA AGG G – 3′ |
| JB_X17 Comp | 5′ – CCT CCG TCC TAG CAA TTG GCT GCT ACC GGA AGG G – 3′ |
| JH | 5′ – CCC TTC CGG TAG CAG CCT GAG CGG TGG TTG AAG G – 3′ |
| JH19 | 5′ – CCC TTC CGG TAG CAG CCT **F**AG CGG TGG TTG AAG G – 3′ |
| JR | 5′ – CCT TCA ACC ACC GCT CAA CTC AAC TGC AGT CTG G – 3′ |
| JR_X8_Comp | 5′ – CCT TCA ACC ACC GCT CAA CTC AAC TTC ATT CTG G – 3′ |
| JR_X17 Comp | 5′ – CCT TCA ACC ACC GCT CAC ATC AAC TGC AGT CTG G – 3′ |
| JH_Duplex_Comp | 5′ – CCT TCA ACC ACC GCT CAG GCT GCT ACC GGA AGG G – 3′ |
| JR_Duplex_Comp | 5′ – CCA GAC TGC AGT TGA GTT GAG CGG TGG TTG AAG G – 3′ |
| JB_Duplex_Comp | 5′ – CCC TTC CGG TAG CAG CCC CTT GCT AGG ACG GAG G – 3′ |
| JX_Duplex_Comp | 5′ – CCT CCG TCC TAG CAA GGA CTC AAC TGC AGT CTG G – 3′ |
| X8_HD | 5′ – TAT GCA GTC ACT ATF AAT CAA CTA CTT AGA TGG T – 3′ |
| X8_HD_Comp | 5′ – ACC ATC TAA GTA GTT GAT TCA TAG TGA CTG CAT A – 3′ |

Table S2. Time-resolved fluorescence decay parameters for Holliday Junctions containing 6-MI

| **DNA Substrate**[1] | $\boldsymbol{\alpha_1}$ [2] | $\boldsymbol{\tau_1}$ | $\boldsymbol{\alpha_2}$ [1] | $\boldsymbol{\tau_2}$ | $\boldsymbol{\alpha_3}$ [1] | $\boldsymbol{\tau_3}$ | $\boldsymbol{\tau_f}$ [2] |
|---|---|---|---|---|---|---|---|
| X11_J3 | 0.47 | 0.63 | 0.48 | 1.59 | 0.05 | 6.89 | 2.75 |
| B12 J3 | 0.77 | 0.50 | 0.20 | 1.38 | 0.03 | 6.28 | 2.03 |
| B16_J3 | 0.59 | 0.45 | 0.17 | 2.7 | 0.24 | 7.4 | 5.47 |
| B17_J3 | 0.42 | 0.43 | 0.28 | 2.58 | 0.31 | 6.98 | 5.55 |
| B18_J3 | 0.46 | 0.47 | 0.26 | 2.98 | 0.28 | 7.07 | 5.53 |
| B19_J3 | 0.45 | 0.67 | 0.32 | 3.74 | 0.23 | 7.71 | 5.59 |
| H19_J3 | 0.8 | 0.54 | 0.14 | 1.83 | 0.06 | 5.78 | 2.57 |
| X16_J3 | 0.84 | 0.29 | 0.09 | 1.88 | 0.07 | 6.11 | 3.4 |

[1]Letter and number indicate location of probe in the J3 junction. [2]Relative amplitudes are calculated from the decays as follows: $\alpha_i = \frac{\alpha_i}{\sum_i \alpha_i}$. [3]$\boldsymbol{\tau_f}$ is the intensity-weighted mean lifetime defined as $\tau_f = \frac{\sum_i \alpha_i \tau_i^2}{\sum_i \alpha_i \tau_i}$

Table S3. Time-resolved fluorescence decay parameters for Holliday Junction or duplex DNA containing 6-MI in the same sequence context.

| **DNA Substrate**[1] | $\alpha_1$ [2] | $\tau_1$ | $\alpha_2$ [2] | $\tau_2$ | $\alpha_3$ [2] | $\tau_3$ | $\tau_f$ [3] |
|---|---|---|---|---|---|---|---|
| 6-MI Monomer | - | - | - | - | 1.00 | 6.52 | 6.52 |
| X8_Duplex | - | - | 0.1 | 3.47 | 0.9 | 7.33 | 7.14 |
| X8_J3 | - | - | 0.17 | 2.81 | 0.83 | 6.82 | 6.53 |
| B12_Duplex | 0.85 | 0.53 | 0.14 | 1.48 | 0.02 | 5.84 | 1.39 |
| B12_J3 | 0.77 | 0.50 | 0.20 | 1.38 | 0.03 | 6.28 | 2.03 |
| B16_Duplex | 0.78 | 0.063 | 0.15 | 2.2 | 0.08 | 7.05 | 3.59 |
| B16_J3 | 0.59 | 0.45 | 0.17 | 2.7 | 0.24 | 7.4 | 5.76 |

[1]Letter and number indicate location of probe in either junction or duplex DNA. [2]Relative amplitudes reported as $\alpha_i = \frac{\alpha_i}{\sum_i \alpha_i}$. [3]$\tau_f$ is the intensity-weighted mean lifetime defined as $\tau_f = \frac{\sum_i \alpha_i \tau_i^2}{\sum_i \alpha_i \tau_i}$

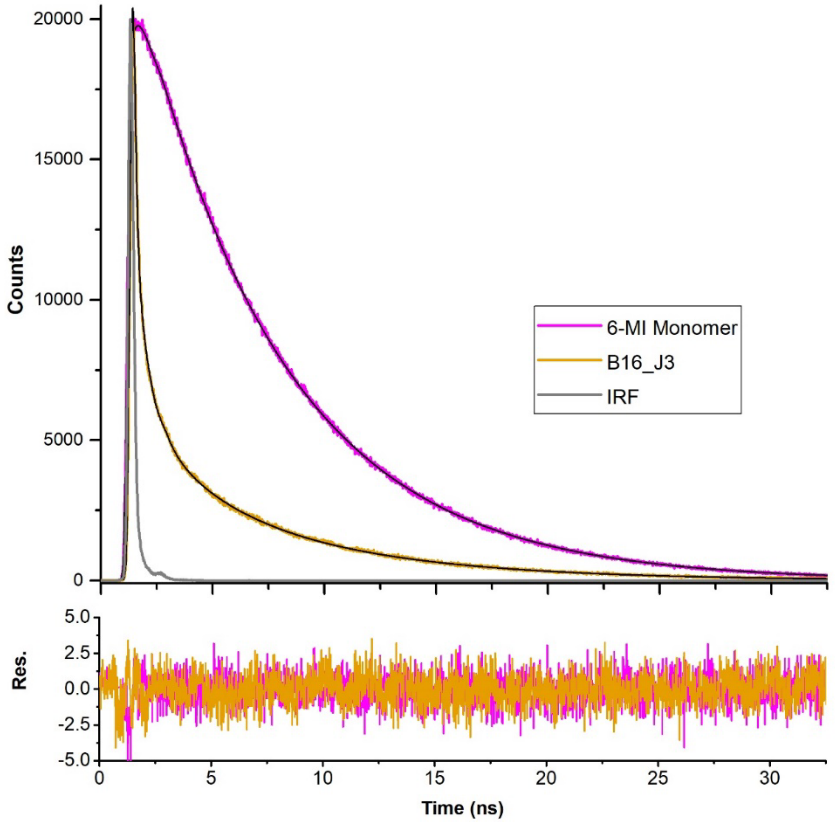

Figure S1. Top: Representative fluorescence lifetime decays of 6-MI monomer (magenta) and B16_J3 (yellow) with fits shown by black lines. The instrument response function is shown in gray. Fluorescence lifetime decays are fit to a sum of exponentials as described in the text. Decays were collected until a maximum of 20000 counts were obtained in the peak channel over a 55 ns time range with 4096 channels. Bottom: The bottom trace depicts the residuals between the fit and the decay. Samples contained 200 nM 6-MI monomer or 6-MI containing DNA in a 10 mM Tris, pH 7.5, 100 mM NaCl and 5 mM $MgCl_2$ buffer. Fit parameters are given in Supporting Information: Table S2.

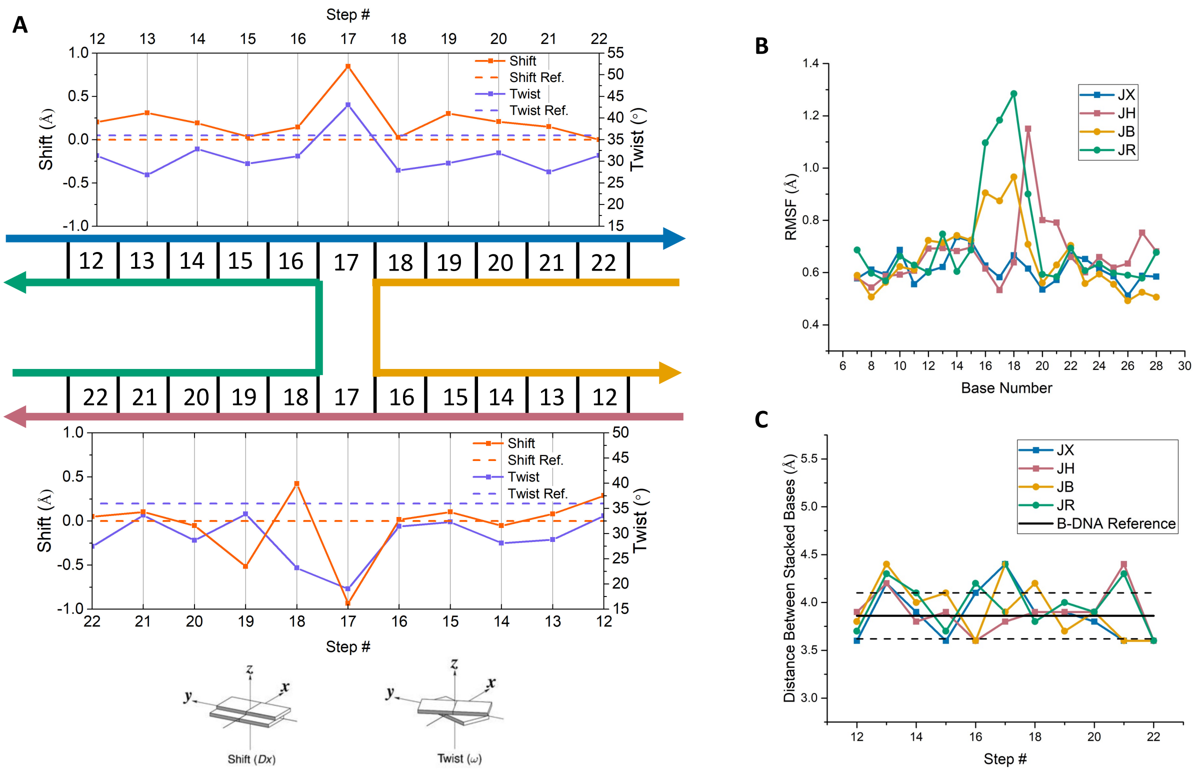

Figure S2. Molecular dynamics simulations of the J3 HJ reveal structural perturbations and increased dynamics at HJ centers. (A) Analyses of a 100 ns simulation were performed using the 3DNA webserver [43] and revealed deviations in the twist (purple) and shift (orange) base pair step parameters of J3 from canonical B-form DNA. Canonical B-DNA values are shown in a dashed line and J3 parameters are depicted with a solid line. Step 17 at the HJ center exhibits the largest deviations. (B) Root-mean-square fluctuations (RMSF) throughout the course of the simulation are shown for bases in the JX (blue), JH (red), JB (yellow) and JR (green) strands. The RMSF values were calculated for individual bases using a sliding window of three bases to estimate the local motion of the middle base with respect to its nearest neighbors. The greatest fluctuations are detected for bases at the center of the HJ in the exchanging strands (JB and JR, yellow and green, respectively). (C) Distances between the center of mass of adjacent bases at each base step in the average MD structure. The distance between bases at the HJ center are greater than the average value determined for B-DNA. The standard deviation for the B-DNA reference is shown by the black dashed lines.

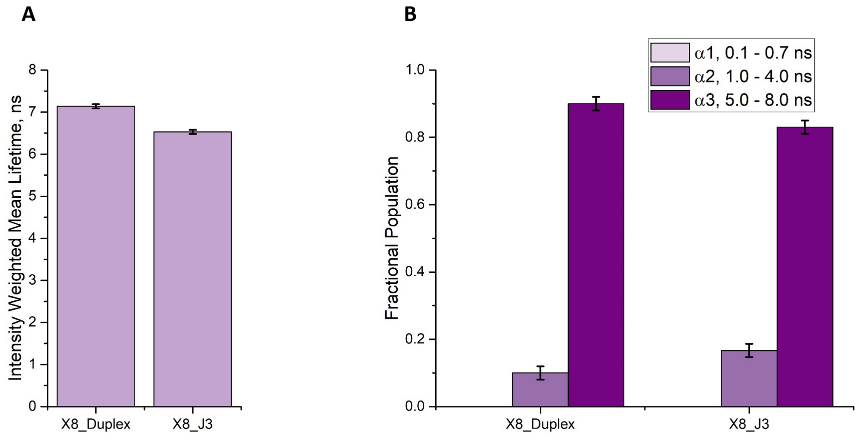

Figure S3. Fluorescence lifetime measurements comparing the DEF sequence in HJ and duplex DNA. (A) Intensity-weighted mean fluorescence lifetimes for 6-MI probes in HJs and duplex DNA within the enhanced fluorescence ATFAA sequence context. (B) Fractional populations of the lifetime components obtained by analyzing fluorescence lifetime decays to a sum of exponentials as described in the text. In this sequence context, shorter lifetimes are observed for the HJ, as the structural constraints of the DEF sequence structure are released, and more dynamic quenching is observed. Samples were 200 nM DNA in a 10 mM Tris pH 7.5, 100 mM NaCl and 5 mM $MgCl_2$ buffer. Fit parameters are given in Supporting Information: Table S3.
